# Spatial transcriptomics deconvolution at single-cell resolution by Redeconve

**DOI:** 10.1101/2022.12.22.521551

**Authors:** Zixiang Zhou, Yunshan Zhong, Zemin Zhang, Xianwen Ren

## Abstract

Computational deconvolution with single-cell RNA sequencing data as reference is pivotal to interpreting spatial transcriptomics data, but the current methods are limited to cell type resolution. Here we present Redeconve, an algorithm to deconvolute spatial transcriptomics data at single-cell resolution, enabling interpretation of spatial transcriptomics data with thousands of nuanced cell states. We benchmarked Redeconve with the state-of-the-art algorithms on diverse spatial transcriptomics datasets and platforms and demonstrated the superiority of Redeconve in terms of accuracy, resolution, robustness, and speed. Applications to a human pancreatic cancer dataset revealed cancer clone-specific T cell infiltration, and application to lymph node samples identified subtle cellular surroundings between IgA+ and IgG+ spots, providing novel insights into tumor immunology and the regulatory mechanisms underlying antibody class switch.

## Main Text

Spatial transcriptomics (ST) technologies provide new tools to identify the cellular organization and interactions of biological samples, which is pivotal to biomedical studies. Multiple ST technologies have been developed and applied to mouse and human brains, heart, lymph node, heart, etc., providing novel insights into cellular communication networks underlying different conditions that were absent before applications of ST. However, ST data are essentially of a spot(pixel)-by-gene matrix structure, needing additional data to provide the cellular information. Integrative analysis of ST data together with matched single-cell RNA sequencing (scRNA-seq) data has provide a transcriptome-wide solution to this question, and multiple effective and efficient algorithms have been proposed.

The current algorithms can be categorized to two groups: (1) mapping-based methods, e.g., NovospaRc^1^, Tangram^2^, Celltrek^3^, and CytoSPACE^4^, which map single cells to the positions of ST data according to gene expression similarity or related measures; and (2) deconvolution-based methods, e.g., CARD^5^, RCTD^6^, cell2location^7^, DestVI^8^, SpatialDWLS^9^, SPOTlight^10^, STRIDE^11^, CellDART^12^, Celloscope^13^, DSTG^14^, and Stereoscope^15^, which try to reconstruct the ST observations by modeling the experimental process as sampling from different combinations of single cells. Mapping-based methods are superior to the current deconvolution-based methods regarding their single-cell resolution as the resolution of current deconvolution methods is limited to tens of cell types. However, mapping-based methods may introduce artificial biases during the mapping process due to the absence of the constraint of reconstruction of the ST observations. It is urgently needed to develop a deconvolution-based algorithm with single-cell resolution to fully release the biological information hidden in ST data.

In this study, we develop a new algorithm, named as Redeconve, to estimate the cellular composition of ST spots. Different from previous deconvolution-based algorithms, Redeconve introduces a regularizing term to solve the collinearity problem of high-resolution deconvolution, with the assumption that similar single cells have similar abundance in ST spots. This algorithmic innovation not only improves the deconvolution resolution from tens of cell types to thousands of single cell states, but also greatly improve the reconstruction accuracy of ST data, enabling illustration of the nuanced biological mechanisms hidden in the ST data. Stringent comparison with the state-of-the-art algorithms including cell2location, CARD, DestVI, CellTrek, NovoSpaRc, and Tangram demonstrates the superiority of Redeconve in terms of reconstruction accuracy, cell abundance estimation per spot, sparseness of the reconstructed cellular composition, cell state resolution, and computational speed. Application to human pancreatic cancer data reveals new insights into tumor-infiltrating CD8+ T cells, and application to human lymph node data reveals new clues for the regulatory factors of IgA+ and IgG+ B cells.

## Results

### Redeconve: a quadratic programming model for single-cell deconvolution of ST data

Redeconve uses scRNA-seq or single-nucleus RNA-seq (snRNA-seq) as reference to estimate the abundance of different cell states in each spot of ST data (Fig. 1a). Different from previous deconvolution methods, Redeconve does not need to group single cells into clusters and then do deconvolution. Instead, Redeconve treats each cell of the sc/snRNA-seq data as a specific cell state serving as reference to estimate the cell composition of ST data. The direct usage of sc/snRNA-seq data as reference is computationally efficient, but will introduce a new challenge, *i.e*., collinearity. That is, multiple single cells have similar profiles of gene expression, prohibiting the accurate estimation of the individual abundance. We introduce a biologically reasonable heuristic by assuming that similar cells have similar abundance within spots of ST data, and thus mathematically introduce a regularization term in the deconvolution model based on non-negative least regression. Solving this regularized deconvolution model by quadratic programming will produce estimation of the cellular composition at single-cell resolution for each spot of ST data.

**Fig 1.**
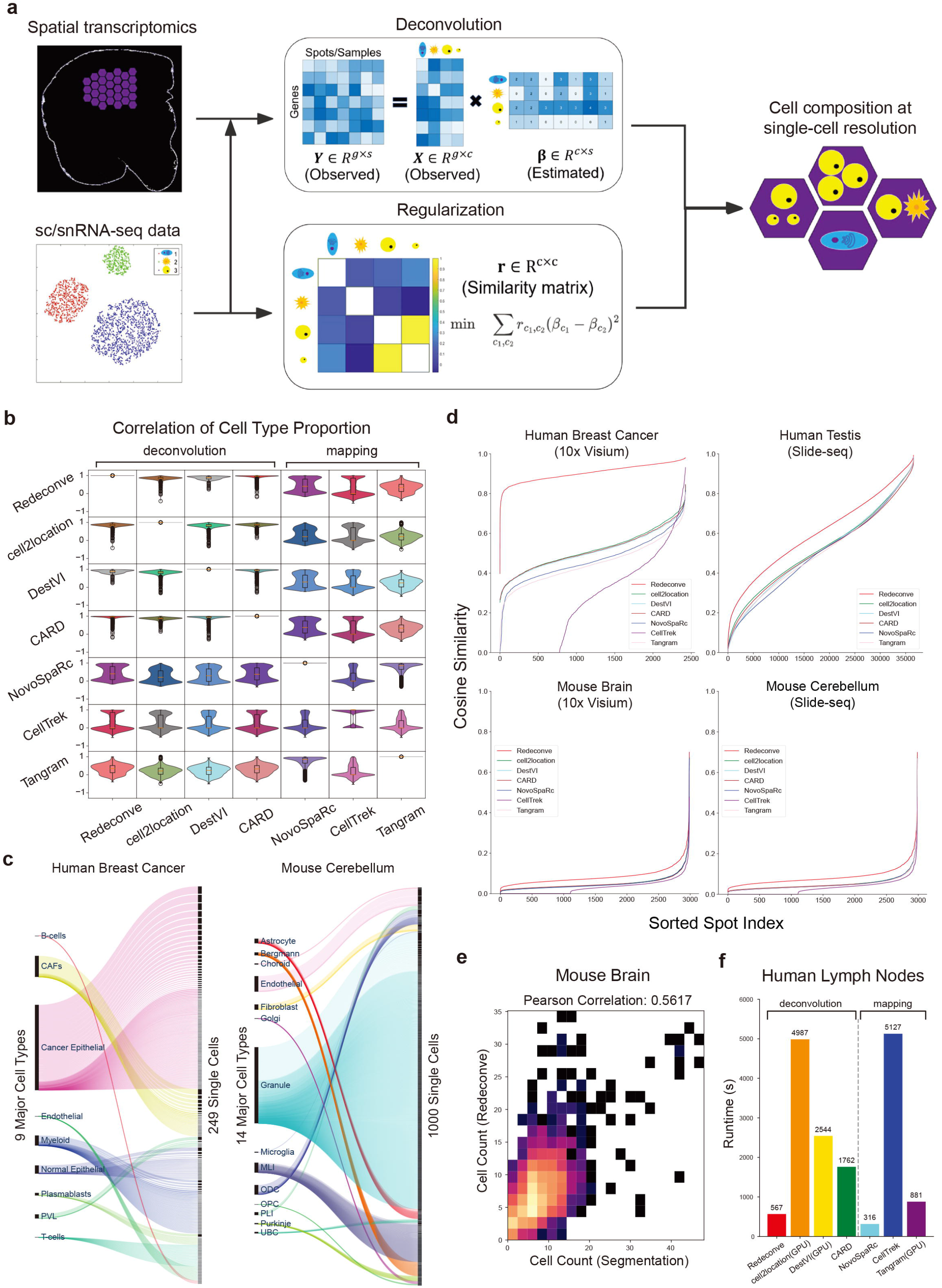
Overview of the Redeconve algorithm and benchmark analysis. a, overview of Redeconve workflow for deconvoluting spatial transcriptomics data. Redeconve requires sc/snRNA-seq data together with spatial transcriptomics data as input and performs deconvolution by solving a regularized non-negative least regression model with the aims to estimate cellular composition across spots at single-cell resolution. b, spot-level pairwise Pearson correlation of cell type proportions among different algorithms on a human breast cancer dataset. c, Sankey diagram demonstrated the cell-type and single-cell resolutions of Redeconve results on human breast cancer and mouse cerebellum datasets, respectively. The bar height of cell types or single cells refer to their estimated abundance after deconvolution. d, line chart of cosine similarities between true and reconstructed expression profiles per spot based on four ST datasets. Spots were sorted by an ascending order of the cosine similarities. e, Pearson correlation of cell abundances between Redeconve and the cell counts per spot based on a mouse brain dataset. The ground truth cell counts per spot was obtained by nucleus counting of cell segmentation image^7^. f, computational efficiency of different deconvolution-based and mapping-based algorithms on a human lymph nodes dataset.

### High accuracy, resolution, robustness, efficacy, and scalability of Redeconve

We applied Redeconve to multiple ST datasets from various platforms (10x Visium, Slide-seq v2, ST, *etc*.) and compared the performance with other methods. We first compared the consistency of results among different methods at the cell type resolution based on a human breast cancer dataset^16^. The results suggested that deconvolution-based methods including Redeconve had higher consistency with each other than mapping-based methods (Fig. 1b), indicating the relative superiority and robustness of deconvolution-based methods. Different from previous deconvolution-based methods which only reported cell-type-level results, Redeconve can further dictate fine-grained cell states at single-cell resolution (Fig. 1c). On a ST dataset from a human breast cancer sample, Redeconve resolved 249 different cell states from 9 major cell types (Fig. 1c). On a ST dataset from mouse cerebellum^17^, Redeconve resolved 1000 different cell states from 14 major cell types (Fig. 1c). In contrast, the resolution of previous deconvolution methods is limited by the clustering results of sc/snRNA-seq data.

In addition to the robustness and resolution superiority, Redeconve further improves the reconstruction accuracy of gene expression per spot (measured by the cosine similarity between the true ST gene expression vector and the reconstructed gene expression profile, Fig. 1d), the accuracy of estimated cell abundance (based on a ground truth by nucleus counting, Fig. 1e and supplementary Fig. 1), and computational speed (Fig. 1f). When suitable reference is provided, e.g., matched scRNA-seq data, Redeconve can reach >0.8 cosine accuracy for most ST spots (Fig. 1d). With no suitable reference available (for example, only snRNA-seq data are accessible for brain samples), Redeconve still outperforms other methods (Fig. 1d). Pairwise comparison between Redeconve and other methods further shows the superiority of Redeconve on almost all spots regarding the reconstruction accuracy (Supplementary Fig. 2-7). Redeconve conducts deconvolution analysis spot by spot, enabling flexible and massive potency for parallel analysis and thus superior computation speed (Fig. 1f).

Redeconve is also superior to estimate the absolute abundance of cells within ST spots. We applied Redeconve to a mouse brain dataset of which the cell counts were obtained by nucleus counting based on image segmentation^7^. Without any priori information, the results of Redeconve showed high conformity with the “ground-truth” cell counts (Fig. 1e), similar to those methods with cell counts (or cell density) as priori knowledge e.g., cell2location and Tangram (Supplementary Fig 1). We used Shannon entropy to estimate the number of different cell states within each spatial spot. Redeconve revealed high spot heterogeneity by showing that some spots had complex cellular composition while others had a relatively simple one. In contrast, the entropy of other methods is uniformly high, showing that each spot was composed of almost all the cell types in reference, which is less realistic (Supplementary Fig. 8).

### Single-cell resolution is unique to Redeconve compared with previous deconvolution algorithms

Then we examined whether the current deconvolution-based algorithms could be upgraded to single-cell resolution by switching the required cell types to thousands of single cells as Redeconve does. Among all the methods we evaluated, only cell2location and DestVI completed the task but took a rather long time compared with the cell type inputs (Supplementary Fig. 9) while other algorithms reported errors. Although single-cell inputs improved the reconstruction accuracy of cell2location on the ST data of a human lymph node sample based on the 10x Genomics Visium platform, cell2location did not reach improvement on the human pancreatic tumor and mouse brain datasets and DestVI failed on all three evaluations (Supplement Fig. 10). In contrast, Redeconve outperformed cell2location and DestVI on almost all spots of the evaluated datasets (Fig. 2a). When switching the inputs from cell types to single cells, DestVI achieved well sparsity regarding the different cell states within each spot (measured by perplexity according to Shannon entropy), similar to the performance of Redeconve. But cell2location reported extremely high perplexity for most spots, indicating high false positive rate (Fig. 2b). Furthermore, we evaluated the computational stability of these three algorithms by repeating all the methods for three times on the same ST data. Redeconve reported identical results because of the deterministic model (Fig. 2c). Cell2location also showed high level of repeatability, but the repeatability of DestVI was not satisfying (Fig. 2c). Therefore, the single-cell resolution and other superiority of Redeconve analysis is mainly derived from algorithmic innovation.

**Fig 2.**
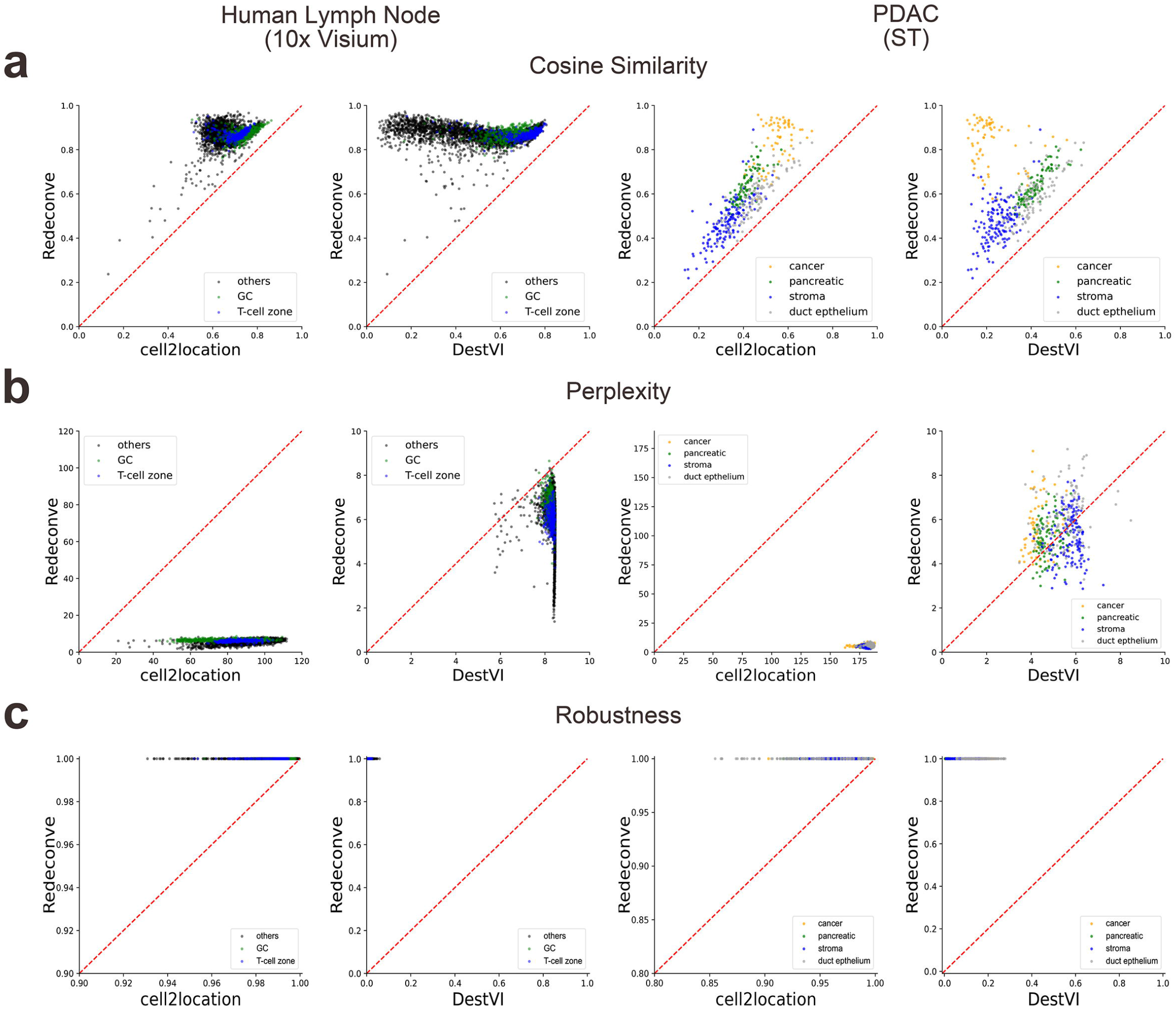
Pairwise comparison of Redeconve with cell2location and DestVI with 1000 single cells as input instead of cell types. Redeconve, cell2location and DestVI are currently the only three deconvolution-based tools with the ability to handle thousands of cell states. a, cosine similarity between true and reconstructed spatial expression profiles. Each dot represents a spot of the ST data. b, the number of different cell states within each spot estimated by the perplexity of cell state composition per spot (See Methods for details). Redeconve and DestVI generated results of tens of cell states, approximating to the true cell numbers per spot while cell2location over-estimated the cellular states of each spot, potentially resulting in false positive estimation. c, reproducibility of different runs. Each method is repeated three times and the average Pearson Correlation Coefficient is calculated among different runs (See Methods for details). Each dot represents a spot. The ST datasets used here were indicated at the top of the figure.

### Single-cell resolution by Redeconve enables identification of pancreatic cancer clone-specific T cell infiltration

To demonstrate the power of deconvolution at single-cell resolution on solving practical biological problems, we deeply analyzed the Redeconve results of the human pancreatic ST dataset^18^. The spatial transcriptomics is from the original ST platform, and scRNA-seq data from the same individual were obtained through InDrop. Histological analysis based on H&E staining annotated four tissue regions: pancreatic, cancer, duct epithelium, and stroma^18^ (Fig. 3a). Redeconve outperformed other methods regarding the reconstruction accuracy for almost all the spots (Fig. 3b-c and Supplementary Fig. 2). Redeconve and CARD showed relatively clear boundaries of tissue regions in accordance with histological analysis (Fig. 3d). Meanwhile, cell2location and DestVI showed blurred boundaries, and NovoSpaRc and Tangram did not show boundaries (Fig. 3d). Further inspection into a specific spot in the upper cancer region (Fig. 3d, the upper zoomed-in piechart) shows that deconvolution-based methods are able to detect fibroblast, which is known to be abundant in pancreatic cancer, while mapping methods fail this task.

**Fig 3.**
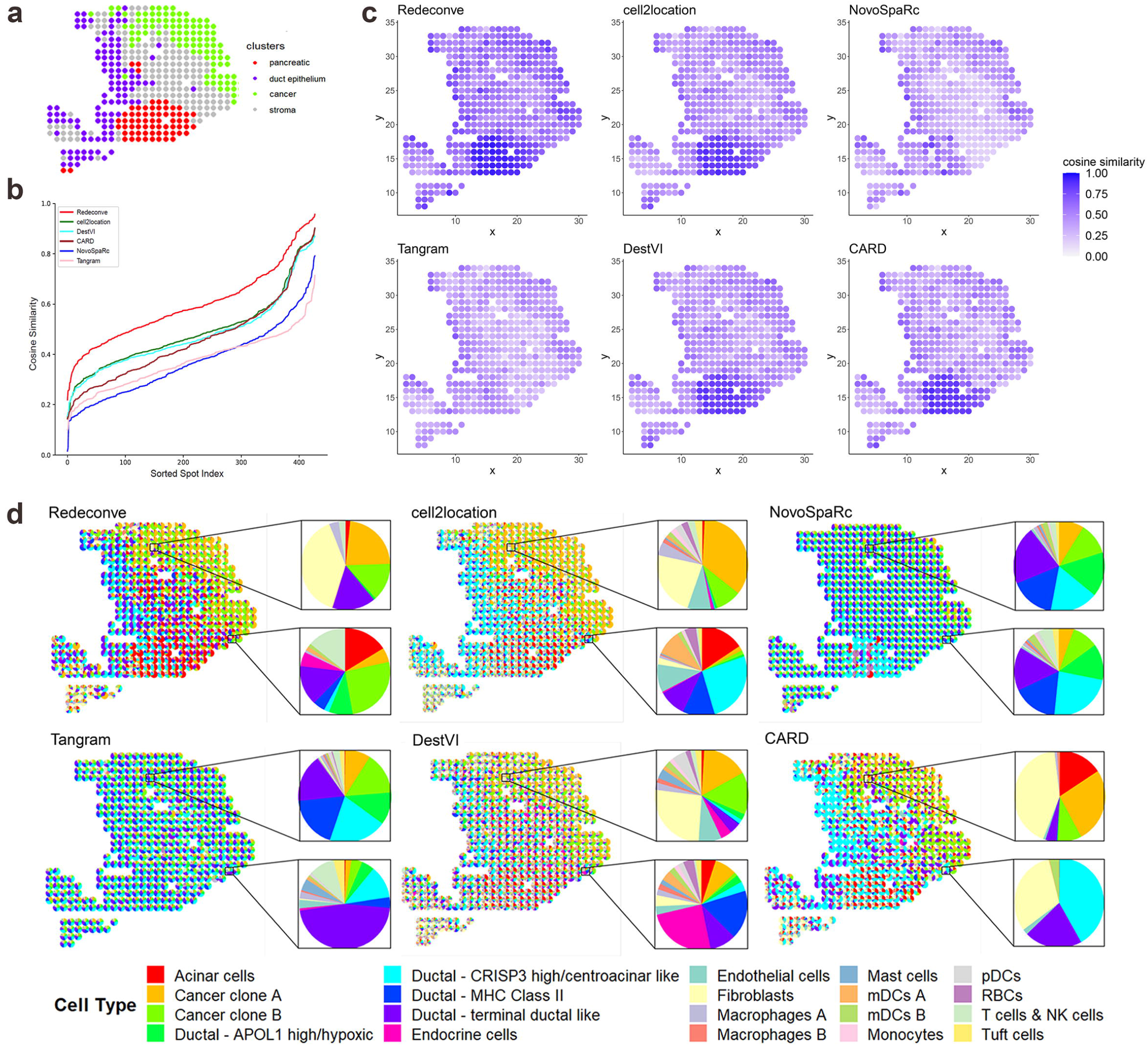
Single-cell deconvolution of a human pancreatic ST dataset. a, four regions were annotated by histological analysis of the original paper: pancreatic, ductal, cancer and stroma regions^22^. b, line chart of cosine similarities between true and reconstructed expression profiles per spot by different computational methods. An ascending order across spots was applied to demonstrate the performance of different methods. c, spatial distribution of the cosine similarity between true and reconstructed expression profiles per spot by different computational methods. d, pie charts displaying the spatial distribution of the estimated cell type proportion per spot by different computational methods.

Then we examined the detailed characteristics of tumor-infiltrating T cells based on these results, which is important to understand the tumor immune microenvironment of pancreatic cancers. The results of cell2location, NovoSpaRc, Tangram and DestVI reported T cells in almost all spots (Fig. 4a), which is not so reasonable in such a cancer tissue. Meanwhile, Redeconve and CARD clearly suggested the sparsity of tumor-infiltrating T cells in pancreatic cancer (Fig. 4a). As CARD is limited by the cell-type resolution, it is difficult to provide more detailed insights. But Redeconve analysis enables deeper investigation. We identified three T cells in the reference scRNA-seq data that appeared in multiple ST spots, indexed as “T.cell.8”, “T.cell. 11” and “T.cell.35” separately (Fig. 4b). By looking up their expression profiles in the reference scRNA-seq, we identified T cell 11 as regulatory T cell (*CD4*^+^ *FOXP3*^+^) and 8 and 35 as *CD8*^+^ cytotoxic T cells. Notably, almost all the T cells within cancer region were similar to regulatory T cell 11, and T cell states similar to 8 and 35 only appeared outside or at the edge of the cancer region (Fig. 4b), indicating the immune suppressive status of the cancer region.

**Fig 4.**
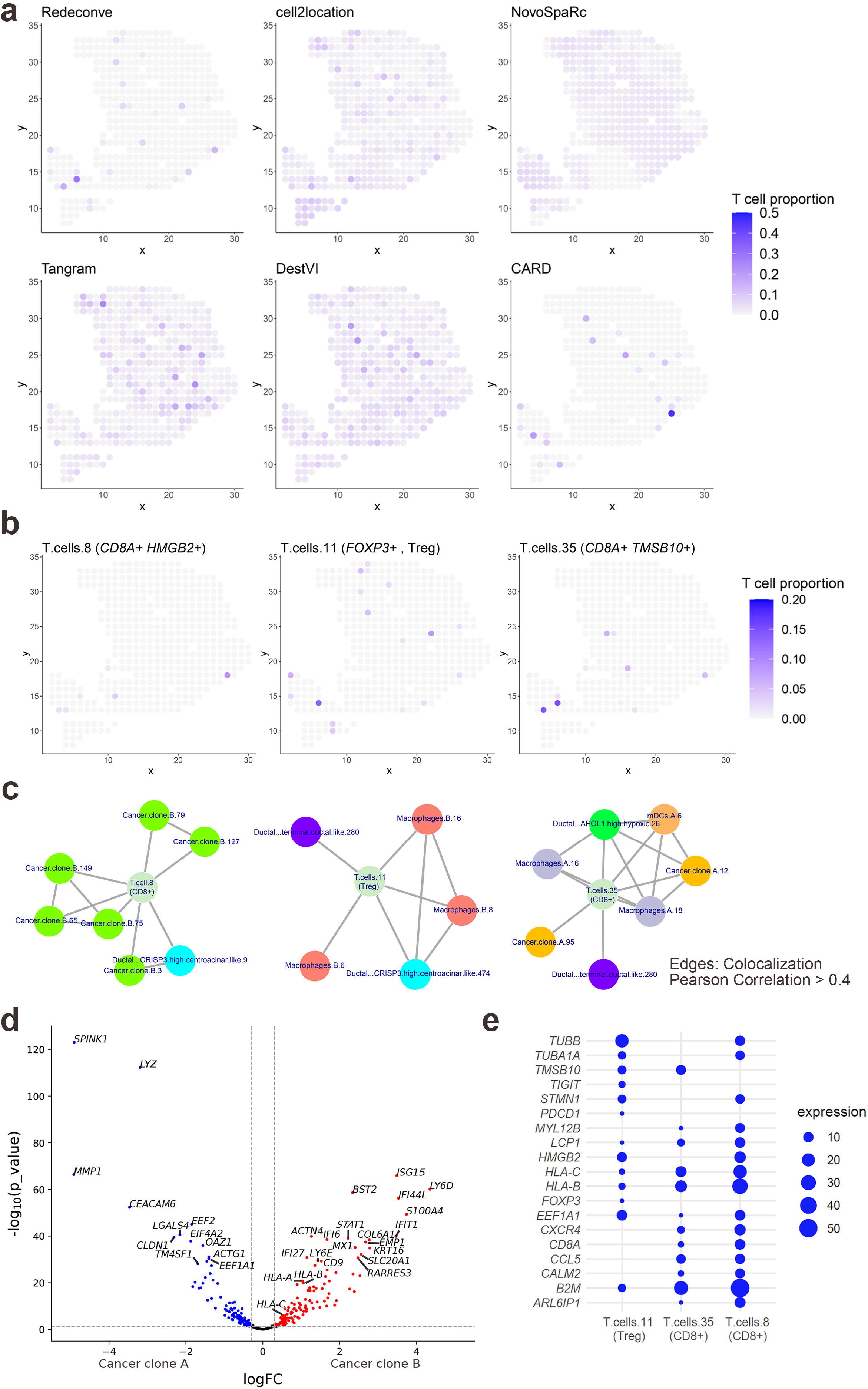
Cancer clone-specific CD8+ T cell infiltration revealed by Redeconve in human pancreatic cancer. a, abundance of T cells per spot estimated by different methods. b, single-cell identity of infiltrated T cells revealed by Redeconve. The three T cells are indexed as “T.cells.8”, “T.cells.11”, “T.cells.35” separately. c, co-localization of the three T cell states with other cellular states. Nodes represent single cells and edges represent co-localization (Pearson correlation of cell abundance > 0.4). Cancer clone-specific CD8+ T cell infiltration was revealed. d, volcano plot displaying differentially expressed genes between the two cancer clones. The blue and red points refer to up-regulated genes in clones A and B-enriched spots, respectively. Vertical dashed line shows the cutoff of log fold change (±0.3). Horizontal dashed line shows the threshold of -lg p (1.301). T cell response-related genes including interferon-stimulating genes and human leukocyte antigens were up-regulated in clone B-enriched cells. e, dot plot displaying characteristics genes among the three T cell states with different spatial preference with cancer clones A and B.

We further conducted co-localization analysis of these three T cell states with the resting cell states by calculating the Pearson correlation coefficient of abundance across all spots (Fig. 4c). The results suggested that the regulatory T cell state similar to T cell 11 mainly co-localized with macrophages similar to macrophages B. 6, 8, and 16 together with duct cells of two different states. Interestingly, T cell 8 and 35 were mainly co-localized with cancer cells, indicating dispersed cancer cells outside the cancer region, which is missed by other methods (Fig. 4c). Furthermore, these two T cell states were separately co-localized with different cancer clones, with T cell state 8 co-localized with cancer clone B and 35 with cancer clone A. Differential gene expression analysis based on the reference scRNA-seq data further indicated the differences between these two pairs of T cells and cancer cells (Fig. 4d-e). It is revealed previously that *TM4SF1*+ cancer cells denoted late-stage while *S1004A*+ cancer cells (clone B) denoted early-stage^19, 20, 21^. Our analysis identified the co-existence of *TM4SF1*+ cancer cells (clone A) and *S1004A*+ cancer cells (clone B) with different *CD8*^+^ T cells, which is important to understand the interactions between cancer and T cells. We found that interferon-induced genes (*IFIT1* and *IFI44L*, for example) and HLA-related genes (*HLA-A*, *HLA-B* and *HLA-C*) were all up-regulated in cancer clone B (Fig. 4d), and correspondingly T cell state 8, which is colocalized with cancer clone B, had high expression of *HMGB2*, *HLA-B* and *HLA-C* (Fig. 4e), indicating well-stimulated T cell response. In contrast, T cell state 35 was *HMGB2*-negative, *HLA*-low and *TMBS10*-positive and co-localized with more A-type macrophages, indicating a less efficacy state. Therefore, with accurate deconvolution at the single-cell resolution, Redeconve can reveal detailed cell-cell interaction at single-cell level and enables discoveries revealing the underlying mechanisms of tumor immunity.

### Redeconve sheds novel insights into the regulatory mechanisms underlying antibody class switch

Redeconve were further applied to analyze an ST data of human secondary lymphoid organs^7^. We again compared Redeconve with other methods on this dataset. In terms of cosine similarity-based reconstruction accuracy, Redeconve achieved mean similarities of 0.868 and significantly outperformed other methods (Fig. 5a). Redeconve achieved high reconstruction accuracy for almost all spots, while, as for other methods, low similarities regions were obvious (Supplementary Fig. 11). We further checked the sparsity of the results by calculating L0-norm. L0-norm of Redeconve has a reasonable distribution between 4 and 32, indicating that only dozens of cell states appear in one spot. In contrast, other methods except CellTrek demonstrated results that almost all cell types appeared in every spot. CellTrek, a mapping-based algorithm, reached low level of L0-norm by generating many “zero-cell” spots, of which Redeconve successfully reconstructed the cellular composition (Supplementary Fig. 12).

**Fig 5.**
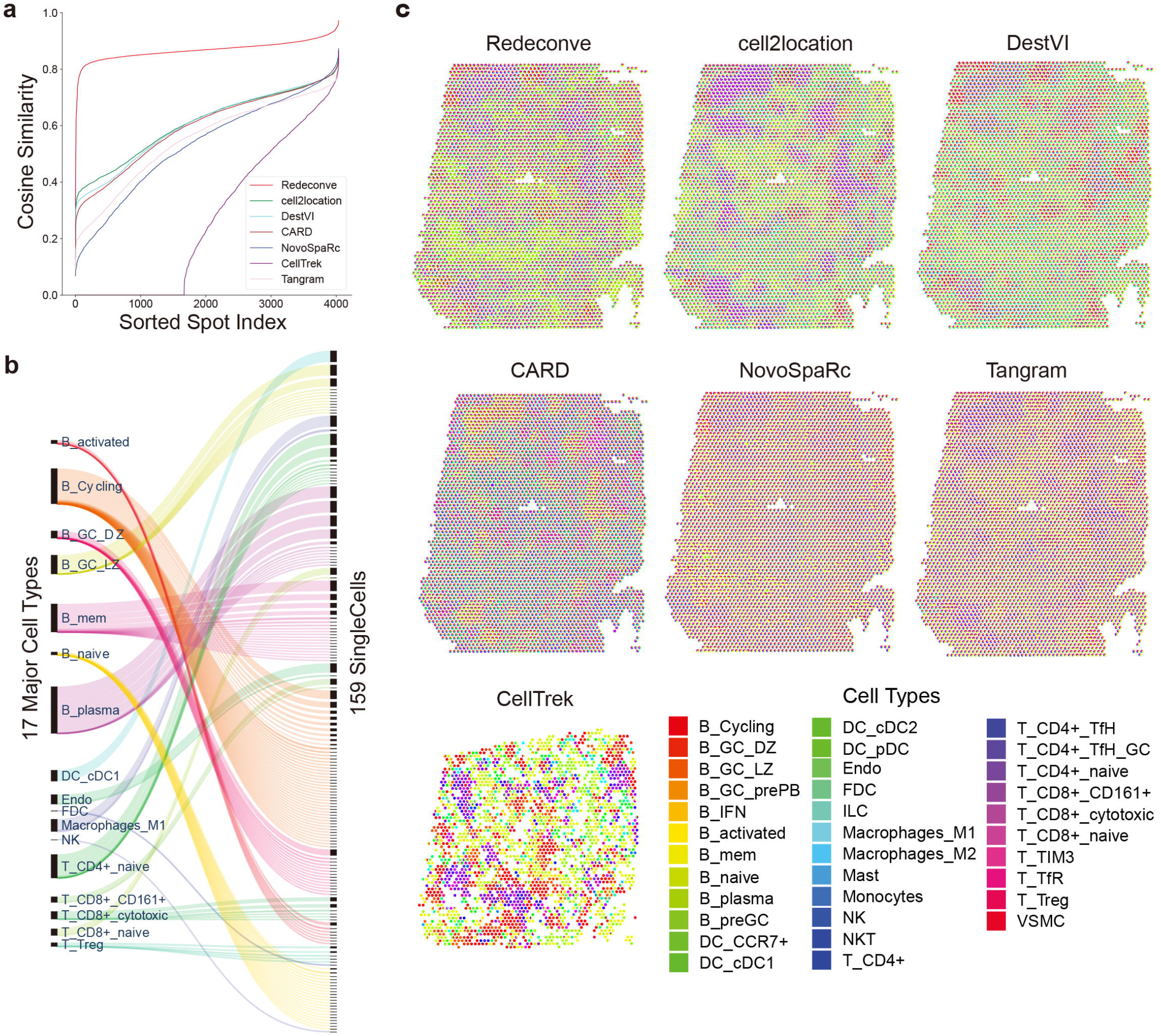
Single-cell deconvolution of a human secondary lymphoid organ ST dataset by Redeconve. a, line chart of cosine similarities between true and reconstructed expression profiles by different algorithms. An ascending order across spots was applied to show the performance of different algorithms. b, Sankey diagram showing the major cell types and single cell states revealed by Redeconve. c, pie chart displaying the spatial distribution of the estimated cell type proportion by different methods.

We further characterized the spatial heterogeneity at single cell resolution to explore the potential regulators of antibody class switch based on this human lymph node data. During the antibody maturation, an activated B cell can change its antibody production from IgM to either IgA, IgG, or IgE depending on the functional requirements, which is termed as class switching. However, the detailed regulators underlying antibody class switching is unclear. Consistent with previous examples, Redeconve outperformed other methods in reconstructing the ST gene expression profiles for almost all spots (Fig. 5a). Spatial pie chart showed that Redeconve produced obvious regional division, while other methods showed blurred or even no boundaries (Fig. 5c). CellTrek failed to analyze some of the spots. Furthermore, compared with cell-type deconvolution, Redeconve identified 159 different cell states from 17 cell types (Fig. 5b). 12 different B plasma cell states were identified in the ST data, which can be further divided into 3 groups (IgA+, IgG+ and negative) based on the expression of *IGHA* and *IGHG* genes. Interestingly, we found that IgA+ and IgG+ B plasma cells are spatially mapped to spots in different regions with little overlap, which means that we could define IgA+ and IgG+ spots based on the abundance of those B plasma cells (Fig. 6a). Next, we took one spot in each of the two regions for detailed inspection at the single-cell resolution. The cell proportion of the two spots shows that *CD8*+ T cells account for a large proportion in the IgA+ spot, suggesting latent interactions between *CD8*+ T cells and IgA+ B plasma cells (Fig. 6b). To confirm the universality of such phenomenon, we conducted differential gene expression analysis between IgA+ and IgG+ spots to identify up-regulated and down-regulated genes (Fig. 6c). As we expected, *IGHA* and *IGHG* were the most differentially-expressed genes; Genes associated with T cells (*TRAC*, *TRBC2*, *CD3D*, *CD8A* for example) were more up-regulated in IgA+ spots, confirming the existence of such interaction. Since lymph node is one of the organs that generate IgA+ plasma cells, the IgA+ spots might be the potential induction sits for IgA+ plasma cells, and CD8+ T cells may play an important role in such process (Fig. 6c). Further co-localization analysis provides more insights (Fig. 6d). We found co-localization of IgA+ plasma cells with CD8+ cytotoxic T cells, suggesting that CD8+ cytotoxic T cells may play important roles during the formation of IgA+ plasma cells. Furthermore, the co-location of IgG+ plasma cells and macrophages may indicate the roles of macrophages during the genesis of IgG+ plasma cells (Fig. 6d). Hence, deconvolution at single cell resolution by Redeconve gains additional insights that may be helpful for uncovering previously opaque biological question.

**Fig 6.**
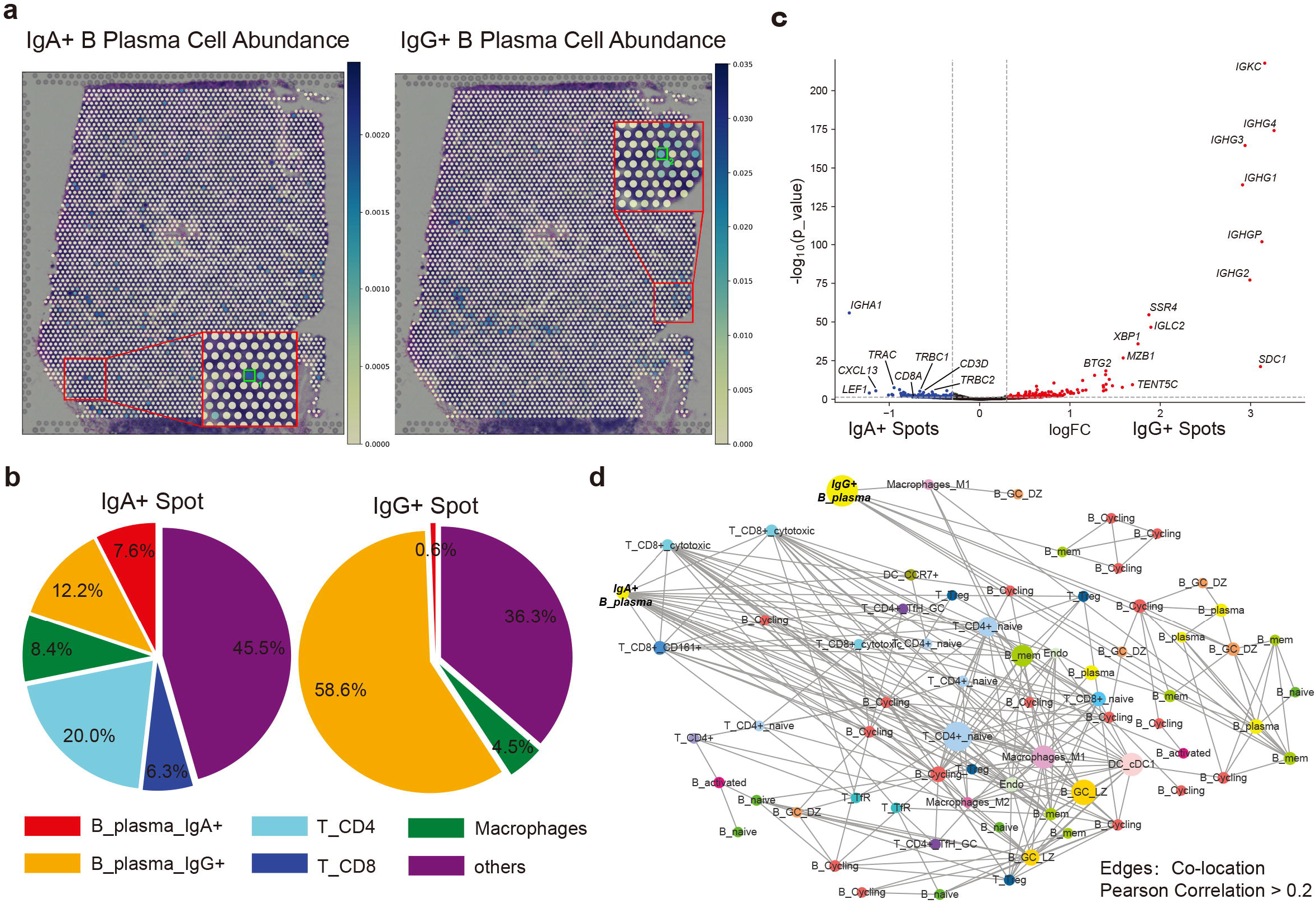
Differences between IgA+ and IgG+ spots regarding cellular composition revealed by Redeconve. a, spatial distribution of IgA+ and IgG+ B plasma cells revealed by Redeconve. b, comparison of the cell proportion of two selected spots (the IgA+ and IgG+ spots in Fig. 6a with green squares). c, volcano plots showing the differential gene expression between IgA+ and IgG+ spots. The red and blue point refer to up-regulated genes in IgG+ and IgA+ spots respectively. Vertical dashed line shows the cutoff of log fold change (±0.3). Horizontal dashed line shows the threshold of -lg(p), namely 1.301. d, co-localization network of IgA+ and IgG+ B plasma cells within the ST data. Nodes represent single cells and edges represent co-located single cells (Pearson correlation of cell abundance > 0.2).

## Discussion

Integrative analysis of disassociated single-cell and *in situ* spatial transcriptomics data is pivotal to construct a comprehensive map of the cellular composition and interactomes of tissues. However, because of technological limitations, current computational methods for integrative analysis of single-cell and spatial transcriptomic data are limited to the cell type resolution. To deep mine the biomedical information hidden in the single-cell and spatial transcriptomics data, here we present Redeconve, a single-cell resolution deconvolution algorithm for integrative analysis of ST data with sc/snRNA-seq data as reference based on a quadratic programming model with regularization of cell-cell similarity, which enables building of comprehensive spatial maps at single-cell resolution for diverse tissues.

We performed stringent evaluation on multiple datasets from a diverse set of ST platforms. The results suggested superiority of Redeconve compared with the state-of-the-art deconvolution-based and mapping-based algorithms in terms of resolution, accuracy, sparsity, robustness, and computational speed. Such improvement from cell-type to single-cell resolution unlocks novel biological discoveries as exemplified by applications in human pancreatic cancer and lymph node samples.

While Redeconve enables deconvolution at single-cell resolution and thus will be a powerful tool for biomedical discoveries, matching between scRNA-seq and ST data appears to be an important factor determining the quality of deconvolution analysis as shown by our evaluation on different tissues (Fig. 1d). Therefore, construction and selection of reference scRNA-seq data according to the specific ST data configuration will be critical in future applications.

Although Redeconve demonstrates superior computational efficacy compared with the state-of-the-art deconvolution algorithms, the single-cell resolution may require extensive computational cost for resolving thousands of cellular states, especially when the cellular throughput of scRNA-seq technologies increases exponentially. Because of the computational complexity of quadratic programming, Redeconve can currently resolve thousands of cellular states based on a standard machine. An enhanced version based on algorithmic innovation or hardware acceleration is needed to handle scRNA-seq datasets of tens of thousands of cellular states.

Deconvolution at single-cell resolution unlocked by Redeconve may also benefit the imputation of ST data with the aid of the rich information in scRNA-seq data. Redeconve has implemented a function to reconstruct the gene expression profiles of individual spots based on the single-cell deconvolution results based on a parsimony principle. The imputed ST data may be more informative to dissect the cellular states of specific tissues.

In summary, we present a novel algorithm named as Redeconve for conducting deconvolution-based analysis of scRNA-seq and ST data at single-cell resolution. The usage of Redeconve is expected to help mapping the cellular architecture at fine granularity across diverse biomedical situations including tumor, immune, development, neurology, and other health and disease conditions. Applications to human pancreatic cancer and lymph nodes showed the potential of Redeconve to bring completely novel insights due to the single-cell resolution unlocked and the superior technical metrics of Redeconve compared to the current state-of-the-art algorithms. We expect Redeconve will be a useful tool to advance the application of scRNA-seq and ST technologies in diverse research disciplines.

## Acknowledgements

This work was supported by Changping Laboratory and the National Natural Science Foundation of China (32022016, 31991171, and 92159305).

## Author contributions

X.R. conceived this study, designed the algorithm, supervised the analysis, and wrote the manuscript. Z.X.Z developed the software, conducted the data analysis, and wrote the manuscript. Y.Z. conducted the data analysis and wrote the manuscript. Z.M.Z provided valuable discussion on the data analysis and wrote the manuscript.

## Figure legends

**Supplementary Fig 1. Pearson correlation of cell abundances estimated by differerent methods and ground truth (estimated by deep learning-based segmentation^7^)**. a, cell2location (5% quantiles) vs segmentation. b, cell2location (Posterior mean) vs segmentation. c, cell2location (95% quantiles) vs segmentation. d, CellTrek vs segmentation. e, Tangram vs segmentation.

**Supplementary Fig 2. Pairwise comparison between Redeconve and other methods regarding the spot-level reconstruction accuracy on PDAC dataset**. a, Redeconve vs cell2location. b, Redeconve vs DestVI. c, Redeconve vs CARD. d, Redeconve vs NovoSpaRc. e, Redeconve vs Tangram.

**Supplementary Fig 3. Pairwise comparison between Redeconve and other methods regarding the spot-level reconstruction accuracy on human lymph nodes dataset**. a, Redeconve vs cell2location. b, Redeconve vs DestVI. c, Redeconve vs CARD. d, Redeconve vs NovoSpaRc. e, Redeconve vs CellTrek. F, Redeconve vs Tangram.

**Supplementary Fig 4. Pairwise comparison between Redeconve and other methods regarding the spot-level reconstruction accuracy on human breast cancer dataset**. a, Redeconve vs cell2location. b, Redeconve vs DestVI. c, Redeconve vs CARD. d, Redeconve vs NovoSpaRc. e, Redeconve vs CellTrek. f, Redeconve vs Tangram.

**Supplementary Fig 5. Pairwise comparison between Redeconve and other methods regarding the spot-level reconstruction accuracy on human testis dataset**. a, Redeconve vs cell2location. b, Redeconve vs DestVI. c, Redeconve vs CARD. d, Redeconve vs NovoSpaRc. e, Redeconve vs Tangram.

**Supplementary Fig 6. Pairwise comparison between Redeconve and other methods regarding the spot-level reconstruction accuracy on mouse brain dataset**. a, Redeconve vs cell2location. b, Redeconve vs DestVI. c, Redeconve vs CARD. d, Redeconve vs NovoSpaRc. e, Redeconve vs CellTrek. f, Redeconve vs Tangram.

**Supplementary Fig 7. Pairwise comparison between Redeconve and other methods regarding the spot-level reconstruction accuracy on mouse cerebellum dataset**. a, Redeconve vs cell2location. b, Redeconve vs DestVI. c, Redeconve vs CARD. d, Redeconve vs NovoSpaRc. e, Redeconve vs Tangram.

**Supplementary Fig 8. Box and violin plots of spot-level information entropy among all algorithms on the six datasets**. At specific spots, higher entropy indicates more complex of cellular composition. Redeconve generated results of which the cellular complexity was similar to the cell numbers per spot of different ST platforms.

**Supplementary Fig 9. Computational cost of cell2location and DestVI when 1000 single cells or cell type references served as inputs to perform deconvolution on the PDAC, human lymph nodes and mouse brain datasets**. SC: 1000 single cells severd as inputs to approach single-cell resolution of deconvolution. CT: cell type references served as inputs for deconvolution. This test was performed on a single NVIDIA A40 card.

**Supplementary Fig 10. Pairwise comparison of spot-level reconstruction accuracy between single-cell (SC) and cell-type (CT) input in cell2location and DestVI on three datasets**. a, the human lymph nodes dataset. b, the PDAC dataset. c, the mouse brain dataset.

**Supplementary Fig 11. Spatial distribution of reconstruction accuracy for spots among different algorithms on the human lymph nodes dataset**. Color represents degree of cosine similarity. a, Redeconve. b, cell2location. c, DestVI. d, CARD. e, NovoSpaRc. f, CellTrek. g, Tangram.

**Supplementary Fig 12. Spatial distribution of the number of cell types with positive values for the spots of the human lymph nodes dataset**. Color represents the number of cell types with positive values. a, Redeconve. b, cell2location. c, DestVI. d, CARD. e, NovoSpaRc. f, CellTrek. g, Tangram.

## Methods

### Algorithm

#### Model overview

In general, we apply an improved linear regression model to deconvolute ST data at single-cell resolution. Given a single-cell (or single-nucleus) expression matrix *X* with dimensions *n*_genes_ × *n*_cells_ and a ST expression matrix *Y* with dimensions *n*_genes_ × *n*_spots_ as input, Redeconve returns a matrix *β* with dimensions *n*_cells_ × *n*_spots_ indicating the estimated number of each cell in each spot. The goal of our model is to optimize the following loss function for each spot separately:

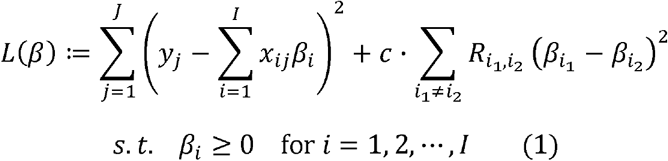

Here *i* = 1,2,···, *I* denotes cells and *j* = 1,2,···, *J* denotes genes. The first term is the traditional Least Square (LS) term and the second term is a regularization term, *c* is a hyperparameter tuning the weight between the two terms. We will later explain the regularization term in details.

Note that this is a typical quadratic programming problem, so we can rewrite our goal as:

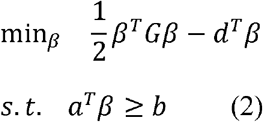

Where *G* is the Hessian matrix, *d*^*T*^ = (2∑_*j*_*y*_*j*_*x*_1*j*_,···, 2∑_*j*_*y*_*j*_*x*_1*j*_), and *a^T^*, *b* are separately

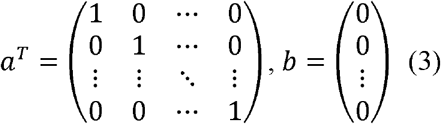

So we can efficiently solve this problem with the *solve*.*QP* function in R package “quadprog” (Turlach B. A. et al. 2013).

#### The regularization term

In sc/snRNA-seq data, the collinearity among cells is serious: cells of the same cell type have very similar expression profiles. This problem would lead to instability of coefficients and reduction of efficiency when directly doing linear regression. To solve this collinearity problem, we further include a regularization term into the loss function. By add this term, we aim at stabilizing the coefficients while having minor effect on the residuals.

In the regularization term *c* ∑_*i*_1_≠*i*_2__ *R*_*i*_1_, *i*_2__ (*β*_*i*_1__ – *β*_*i*_2__)^2^, *R*_*i*_1_,*i*_2__ is a measure of similarity between cell *i*_1_ and *i*_2_, which is

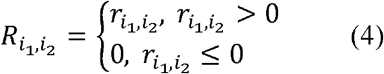

Where *r*_*i*_1_,*i*_2__ is the Pearson correlation coefficient between cell *i*_1_ and *i*_2_. Namely, when the Pearson correlation coefficient is greater than zero, *R*_*i*_1_,*i*_2__ is equal to the Pearson correlation coefficient; otherwise *R*_*i*_1_,*i*_2__ is zero. So, we manually bring the coefficients of cells whose expression profile is similar closer. By doing this, we can guarantee the robustness and precision of our result.

#### Determination of the hyperparameter

A key point of this model is how to select the hyperparameter: an extremely small hyperparameter will make the regularization term ineffective, while an extremely large one will greatly affect the fitting residuals. An ideal hyperparameter should be as large as possible while affecting the fitting residual as little as possible. Here we offer 3 ways to set the hyperparameter:

1. “default”: use the default hyperparameter we set according to the number of cells and genes;
2. “customized”: set the hyperparameter arbitrarily by the user;
3. “autoselection”: automatically calculate and select the optimal hyperparameter.

In mode “default”, we use the following formula to set the hyperparameter:

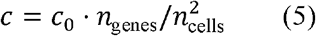

Where *c*_0_ is a predetermined constant and is set to 10^5^. The idea of this formula is: (1) the LS term is approximately proportional to n_genes_, so as n_genes_ increases *c* should synchronously increase; (2) the regularization term is approximately proportional to the square of n_cells_, so as n_cells_ increases *c* should decrease by 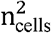.

In mode “autoselection”, we apply the following method to determine the optimal hyperparameter:

1. We first calculate a hyperparameter *c_d_* according to the formula in mode “default”, and set up a series of hyperparameter *c*_1_, *c*_2_, *c*_3_, *c*_4_, *c*_5_ as 0.01*c_d_*, 0.1*c_d_, c_d_*, 10*c_d_*, 100*c_d_*;
2. Then we run deconvolution with these hyperparameters separately, and calculate the residual *ε_i_* for each *c_i_*;
3. We further calculate 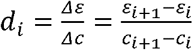;
4. We check these *d_*i*_*, then choose *c_i_* that maximizes *d_i_* as the optimal hyperparameter (This indicates: if the parameter continues to increase, the residual will increase significantly). Namely, we choose *c_i_* that satisfies:

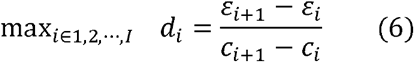

By this procedure, we can get the hyperparameter that maximizes the power of regularization term while having minor effect on the LS term.

### Data preprocessing

To run the deconvolution, the following data preprocessing steps are necessary. Note that some steps are alternative according to users’ needs.

1. Get the expression profiles of cell type/Sampling of single cells. If a cell-type deconvolution is to be run, we will estimate the expression profile *x_ij_* of cell type *i* and gene *j* as the average expression of gene *j* across all cells within cell type *i*. If a single-cell deconvolution is to be run and the number of single cells is overwhelming, we will take stratified samples of cells by cell type to get a rational number of cells.
2. Gene filtering. Deconvoluting with tens of thousands of genes is time-consuming or even misleading, so we select highly variable genes before deconvolution for computational efficacy. Filtering criteria include the following three standards: (1) These genes appear in both sc/snRNA-seq data and spatial transcriptomics; (2) The variance of these genes in sc/snRNA-seq data must be larger than a threshold (default is 0.025); (3) The average counts per spot must be bigger than a threshold (default is 0.003). This finally results in ~8000 genes for deconvolution. Redeconve allows deconvolution without gene filtering with higher computational cost.
3. Normalization of reference. We add a pseudo-count of 0.5 to the “zeros” in sc/snRNA-seq data, and normalize sc/snRNA-seq data to TPM (transcripts per million). Preprocessing operations are not needed for spatial transcriptomics data.

### Datasets for benchmarking

#### PDAC

ST data of a human pancreatic ductal adenocarcinomas (PDAC-A) with 438 spots and sample-matched scRNA-seq data (InDrop) with 1,926 single cells across 20 cell types were integrated by Moncada et al.^22^, and an intersection of 19,736 genes was used in our study. The annotation of four main structural regions based on histological analysis by Moncada et al.^22^ was used during our analysis to depict the spatial characteristics of the ST data.

#### Human lymph node

Human lymph node Visium data were downloaded from the 10x Genomics website (https://www.10xgenomics.com/resources/datasets/human-lymph-node-1-standard-1-1-0), which includes a total number of 4,035 spots. ScRNA-seq data were collected from Kleshchevnikov et al^7^, of which 73,260 cells across 34 cell types were collected. Since this scRNA-seq dataset captured a wide spectrum of immune cell states spanning lymph nodes, tonsils and spleen, we used it as reference to reveal the phenotypic diversity of immune cells when deconvoluting at single cell resolution.

#### Mouse cerebellum

The DropViz scRNA-seq dataset were generated by Saunders A. et al.^17^ and were collected by Cable D. M. et al.^6^ along with the annotations of the cells. This scRNA-seq dataset was processed by Cable D. M. et al.^6^ so that each cell type contains no more than 1,000 cells. The Slide-seq mouse cerebellum data with 39,496 spatial spots were collected by Cable D. M. et al. using the Slide-seq v2 protocol^6^. Both of these datasets were downloaded from https://singlecell.broadinstitute.org/single_cell/study/SCP948/robust-decomposition-of-cell-type-mixtures-in-spatial-transcriptomics#study-download.

#### Human Brest Cancer

Human Brest Cancer Visium data related to the Wu et al. study^16^ was available at https://zenodo.org/record/4739739#.Ys0v6jdBy3D. Sample ‘CID4290’ that includes 2,426 in tissue spots was used for deconvolution. ScRNA-seq data that includes 100,064 single cells with annotations (Access number: GSE176078, the NCBI GEO database) served as reference to do deconvolution analysis.

#### Human Testis

The processed Human Testis Slide-seq dataset was download from https://www.dropbox.com/s/q5djhy006dq1yhw/Human.7z?dl=0 and sample ‘Puck5’ with 36,591 spots was used for evaluation in this study^23^. The reference scRNA-seq data that includes 6,490 single cells was obtained from the NCBI GEO database with access number GSE112013, and the corresponding annotations were available in the supplementary information table S1 by Guo et al^24^.

#### Mouse Brain

10x Visium and snRNA-seq data (includes annotation) were available in the ArrayExpress database with accession numbers E-MTAB-11114 and E-MTAB-11115, respectively^7^. Sample ‘ST8059048’ containing 2,987 spots was used for evaluation in this study, and all 40,532 single cells across 59 cell types served as reference. In addition, the corresponding data of nuclei counts estimated by histological image segmentation based on deep learning s was downloaded from https://github.com/vitkl/cell2location_paper/blob/master/notebooks/selected_results/mouse_visium_snrna/segmentation/144600.csv.

### Comparing Redeconve with alternative methods

We compared Redeconve with recently developed deconvolution-based methods (cell2location^7^, DestVI^8^ and CARD^5^) as well as mapping-based methods (NovoSpaRc^1^, CellTrek^3^ and Tangram^2^).

#### Parameter setting

Prediction results for the 6 datasets were obtained by running the corresponding programs of the algorithms aforementioned based on the default settings except some special considerations: (1) 1,000 cells were randomly selected in NovoSpacRc to avoid large number of total cells; (2) 1,000 stratified samples of cells were used for Redeconve in almost all the datasets except PDAC where we used total 1,926 cells; (3) minCountGene and minCountSpot of the createCARDObject function were set to 0 to prevent unexpected gene or spot filtering in CARD. The output of each method was either a cell-by-spot matrix represented absolute abundance (Redeconve, Tangram) or proportion (NovoSpaRc) of single cells existing at each spot or estimated cell-type abundance (cell2location) or proportion (DestVI, CARD) matrix except CellTrek, of which the outcome was predicted spatial coordinates for individual cells. Hence, for CellTrek, we obtained cell-by-spot abundance matrix by assigning single cells to specific spots according to whether the spot area designed by ST platforms covered the predicted coordinates. We only evaluated CellTrek on the two 10x Genomics Visium-based datasets (human lymph node, human breast cancer and mouse brain) because of running errors on other ST datasets in our computational environment.

#### Calculating Performance Metrics

For all datasets, spot-wise cosine similarities between observed and predicted spot-by-gene expression matrix were calculated. Based on the output of each algorithm, we first calculated the predicted expression matrix through two ways: (1) for Redeconve, NovoSpaRc, CellTrek and Tangram, we multiplied spot-by-cell abundance or proportion matrix by the cell-by-gene sc/snRNA expression matrix; (2) for cell2location, DestVI and CARD, we multiplied the cell-type abundance or proportion matrix by the reference cell-type expression matrix, where the reference was generated through averaging sc/snRNA expression data according to cell types. Then, to estimate sparsity of the results, we calculated cell-type proportion matrices of all programs and then compared them according to cell-type information entropy and L_0_ norm. The L_0_-norm represents number of cell types present at each spot (nonzero values). We also evaluated the performance of cell abundance estimation by Pearson’s correlation between results of individual methods (Redeconve, cell2location, CellTrek and Tangram) and the cell numbers estimated by histological image segmentation based on deep learning for the mouse brain dataset. Finally, computational efficiencies were estimated through comparing total time spent by each algorithm on a computer with Intel(R) Xeon(R) Platinum 8253 CPU, where we set the maximum number of cores to 96. In addition, we tested the run time of these programs on a single NVIDIA A40 card if GPU acceleration supported (cell2location, DestVI, and Tangram).

#### Assessment at Single Cell Resolution

Cell-by-spot abundance matrix is required for comparison among deconvolution-based methods at single-cell resolution. We, therefore, applied Redeconve with 1,926 and 1,000 single cells sampled from the reference scRNA-seq data for the two ST datasets (PDAC and human lymph node), respectively and assigned every single cell a unique cell type since cell2location, DestVI and CARD only support cell-type deconvolution. The result matrices of Redeconve, cell2location and DestVI (no result was available for CARD because of running errors) was obtained according to the corresponding default settings. Cosine similarity, information entropy, perplexity and runtime efficiencies were evaluated as mentioned above. We also estimated the result robustness by repeating three times for each algorithm with the same parameter setting and input and then calculating Pearson’s correlation of single cell abundance among the three repeated runs at each spot. In these cases, global random seeds were set as 0, 1 and 2 for the repeated runs.

#### Information entropy and perplexity

Then we calculate Information entropy *H* and perplexity *P* for each spot separately by the following formula:

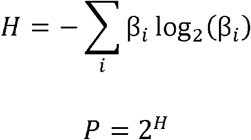

where *i* = 1,2,…, *I* denotes different cell states. *β_i_* were normalized in advance so that their sum equaled to 1 (*i.e*. they denote proportion rather than absolute abundance). Here, the perplexity can approximately denote the number of cell states that occurs in a spot, so we used this index to evaluate the sparsity of each method.

### Downstream Analyses after Redeconve deconvolution

#### Human lymph node

We firstly ran Redeconve on default setting to obtain deconvolution result at single-cell resolution. Then, we investigated the spatial distribution of plasma cells after grouping these plasma cells into IgA+, IgG+ and others based on the expression of *IGHA1*, *IGHG1*, *IGHG3* and *IGHG4*. IgA+ and IgG+ spots were determined by the following three steps: (1) identifying the top 50% spots with the highest abundance of IgA+ and IgG+ plasma cell enriched, which were named as spot sets A and G; (2) identifying the difference sets between A and G, and naming as AD and GD; (3) selecting spots from AD and GD with the top 1% IgA+ and IgG+ plasma abundance, which were assumed to be IgA+ and IgG+ spots respectively. EdgeR^25^ was applied to perform differential gene expression analysis and identified significantly differential genes between IgA+ and IgG+ spots. Then, we calculated Pearson’s correlation coefficient among single cell states in the reference across IgA+ and IgG+ spots and took single cells as nodes and correlated cells (Pearson > 0.2) as edges to generate the cell-cell co-location network.

#### PDAC

We ran Redeconve with all the 1,926 single cells as reference, and all the parameters were kept default. For downstream analyses, we first compared Redeconve with existing tools as described in the aforementioned sections. Then, to study the distribution of T cells, we distinguished from NK cells T cells by the expression of *CD3D*, *CD3E* or *CD3G* in the scRNA-seq data. We further picked out those T cells that frequently appeared in the ST spots (T cells 8, 11, and 35). To study the spatial colocalization of these T cells with other cells, we calculated the Pearson’s correlation of cell abundance across spatial spots, and generated a colocalization network of single cell resolution using those cell pairs whose Pearson correlation were greater than 0.4 with the R package igraph^26^.

## Data availability

### PDAC

Spatial transcriptomics data, scRNA-seq data and the annotation of four main structural regions are all available on Gene Expression Omnibus with accession number <u>GSE111672</u>. The file of the ST data we used was “GSM3036911_PDAC-A-ST1-filtered.txt.gz”, and the file of the scRNA-seq data we used was “GSE111672_PDAC-A-indrop-filtered-expMat.txt.gz”.

### Human lymph node

The spatial transcriptomics (Visium) data were downloaded from the 10x Genomics website (https://www.10xgenomics.com/resources/datasets/human-lymph-node-1-standard-1-1-0). ScRNA-seq data were collected from Kleshchevnikov et al.^7^.

### Mouse cerebellum

Both of spatial transcriptomics and scRNA-seq data are downloaded from https://singlecell.broadinstitute.org/single_cell/study/SCP948/robust-decomposition-of-cell-type-mixtures-in-spatial-transcriptomics#study-download. The spatial transcriptomics data file was “Cerebellum_MappedDGEForR.csv”, and the reference scRNA-seq file was “1000cellsSubsampled_cerebellum_singlecell.RDS.zip”.

### Human Brest Cancer

The spatial transcriptomics (Visium) was available at https://zenodo.org/record/4739739#.Ys0v6jdBy3D (Sample ‘CID4290’). The scRNA-seq data was available on the NCBI GEO database with accession number <u>GSE176078</u>.

### Human Testis

The processed Spatial transcriptomics (Slide-seq) dataset was download from https://www.dropbox.com/s/q5djhy006dq1yhw/Human.7z?dl=0 (Puck5). The scRNA-seq data was available on the NCBI GEO database with accession number <u>GSE112013</u>.

### Mouse Brain

The spatial transcriptomics (10x Visium) and snRNA-seq data (includes annotation) were available in ArrayExpress database with accession numbers <u>E-MTAB-11114</u> and <u>E-MTAB-11115</u>, respectively. We used the ST data of sample ST8059048. Nuclei counts estimated by segmenting histology images was available in https://github.com/vitkl/cell2location_paper/blob/master/notebooks/selected_results/mouse_visium_snrna/segmentation/144600.csv.

## Code availability

Redeconve and the codes used to generate all the pictures in this paper is available at https://codeocean.com/capsule/5481250/tree and https://github.com/ZxZhou4150/Redeconve.

